# EEG-based Graph Neural Network Classification of Alzheimer’s Disease: An Empirical Evaluation of Functional Connectivity Methods

**DOI:** 10.1101/2022.06.14.496080

**Authors:** Dominik Klepl, Fei He, Min Wu, Daniel J. Blackburn, Ptolemaios G. Sarrigiannis

## Abstract

Alzheimer’s disease (AD) is the leading form of dementia worldwide. AD disrupts neuronal pathways and thus is commonly viewed as a network disorder. Many studies demonstrate the power of functional connectivity (FC) graph-based biomarkers for automated diagnosis of AD using electroencephalography (EEG). However, various FC measures are commonly utilised, as each aims to quantify a unique aspect of brain coupling. Graph neural networks (GNN) provide a powerful framework for learning on graphs. While a growing number of studies use GNN to classify EEG brain graphs, it is unclear which method should be utilised to estimate the brain graph. We use eight FC measures to estimate FC brain graphs from sensor-level EEG signals. GNN models are trained in order to compare the performance of the selected FC measures. Additionally, three baseline models based on literature are trained for comparison. We show that GNN models perform significantly better than the other baseline models. Moreover, using FC measures to estimate brain graphs improves the performance of GNN compared to models trained using a fixed graph based on the spatial distance between the EEG sensors. However, no FC measure performs consistently better than the other measures. The best GNN reaches 0.984 area under sensitivity-specificity curve (AUC) and 92% accuracy, whereas the best baseline model, a convolutional neural network, has 0.924 AUC and 84.7% accuracy.

## I. INTRODUCTION

Alzheimer’s disease (AD), a neurodegenerative disease, is the most common form of dementia. AD patients exhibit progressive deterioration of memory and other cognitive functions. From a neuroscience perspective, AD leads to synaptic loss and cellular death, which progressively occurs over multiple brain regions [1]. Disruption of communication pathways amongst brain regions is observed in AD [2–4]. Due to this distributed nature of AD, it can be recognised as a network disorder. Thus, graph theory is well suited for analysing and classifying AD, as it provides a general framework to study the interactions of various pathological processes across multiple spatiotemporal scales.

Functional connectivity (FC) is one of the methods to construct and study brain graphs. The edges of FC graphs represent the statistical dependencies between brain regions rather than the physical connectome, i.e. structural connectivity. FC brain graphs can be constructed from any functional brain imaging modality, such as M/EEG, fMRI, or PET. In this paper, we focus on EEG. EEG has been shown to be an effective tool for studying the changes in brain activity in AD cases [5–7]. Compared to other modalities, EEG is economical, non-invasive, easy to administer, and has a superior temporal resolution. On the other hand, it suffers from a low spatial resolution as the activity is measured by means of electrodes placed on the scalp of the subject.

Emerging evidence shows large-scale alterations in functional connectivity (FC) in AD, such as increased connectivity in the low-frequency bands [8–10]. Graphbased studies show that AD is characterised by reduced complexity [6] and loss of small-world organisation, assessed by the clustering coefficient and characteristic path length [11–14].

However, there are multiple FC methods commonly used within the literature. Furthermore, each FC method may quantify a different aspect of brain coupling. There are both linear and nonlinear FC methods that quantify the coupling of signals’ phases, amplitudes, and even cross-frequency interactions. For simplicity, we do not consider cross-frequency interactions and focus on FC measures quantifying brain coupling between two signals within the same frequency range, i.e. frequency band. Moreover, there is an uncertainty on the reproducibility of some FC methods [15] and the robustness to volume conduction effects [16]. In this paper, we select and evaluate a number of commonly used methods to quantify FC from EEG data.

The most common and simplest FC measure is Pearson’s correlation coefficient (corr) [17]. The phase coupling of a pair of signals is commonly measured by coherence (coh) [13, 14], imaginary part of coherency (iCOH) [15, 18], phase lagged index (PLI) and its weighted version (wPLI), and phase locking value (PLV) [15]. The correlation of amplitude envelopes of two signals can also be measured (amp-corr) [15]. Finally, information-theoretic measures are commonly used as measure FC, such as mutual information (MI) [19]. We exclude the directed (i.e. causal) connectivity measures from this comparative study, since these are generally not considered as FC measures but rather effective connectivity measures.

Graph-based features were also successfully used to train machine learning classifiers to diagnose brain disorders using EEG automatically. Manually engineered graph features, such as node strength [20] and vectorised adjacency matrix [21], could be promising graph-based biomarkers of AD, as both approaches achieve high classification accuracy. Additionally, there were some attempts to utilise deep learning, convolutional neural network (CNN), for automatic graph-based feature extraction. Specifically, CNN was trained to classify AD and schizophrenia using adjacency matrices, which are image-like representations of FC graphs [22]. However, an image representation of a graph cannot effectively capture all the properties, as a graph is a non-euclidean object.

Graph neural network (GNN) extends the logic of convolution operation to graphs by aggregating information from linked nodes, based on the assumption that nodes connected by an edge are similar. However, there is a limited number of GNN applications for EEG brain graph classification. Moreover, it is unclear which method should be used to infer the graph structure for GNN application. A fully connected graph is commonly used in the literature [23]. However, such an approach does not leverage any information encoded by FC brain graphs. A second option is using the distances between spatial positions of EEG electrodes to define the graph structure [23, 24]. Furthermore, Demir et al. [23] utilise distance thresholding and k-nearest-neighbours methods to filter-out unimportant edges. Such edge-filtering can be important, as some edges might be redundant or even introduce additional noise, thus hindering the model from learning the optimal solution. Only a handful of studies use FC measures, such as coherence [24] and wPLI [25]. Additionally, Liu et al. [25] use a minimum spanning tree algorithm (MST) in order to produce sparse brain graphs. This is in contrast to threshold-based edge filtering, as MST can select edges with various edge weights and ensures that the resulting graph is connected. Additionally, Zhong et al. [26] utilise a learnable mask in order to learn the optimal graph structure for a specific classification task without relying on any FC measure.

In this study, we systematically evaluate the effects of using various FC methods to infer EEG brain graphs in training GNN for the classification of AD patients. Two types of edge filtering are used to induce graph-sparsity in order to improve the performance of GNN. To compare and evaluate the classification performance of various FC-based GNNs, a GNN-based baseline is trained using a fixed graph structure for all brain graphs, represented by the euclidean distance between spatial positions of EEG sensors. Three additional baseline models are established: two SVM baselines fitted on node strength (SVM-NS) and vectorised adjacency matrix (SVM-vector), respectively, and a CNN trained on images of adjacency matrices. Fig. 1 illustrates the model architectures employed for comparative study in this work.

**FIG. 1:**
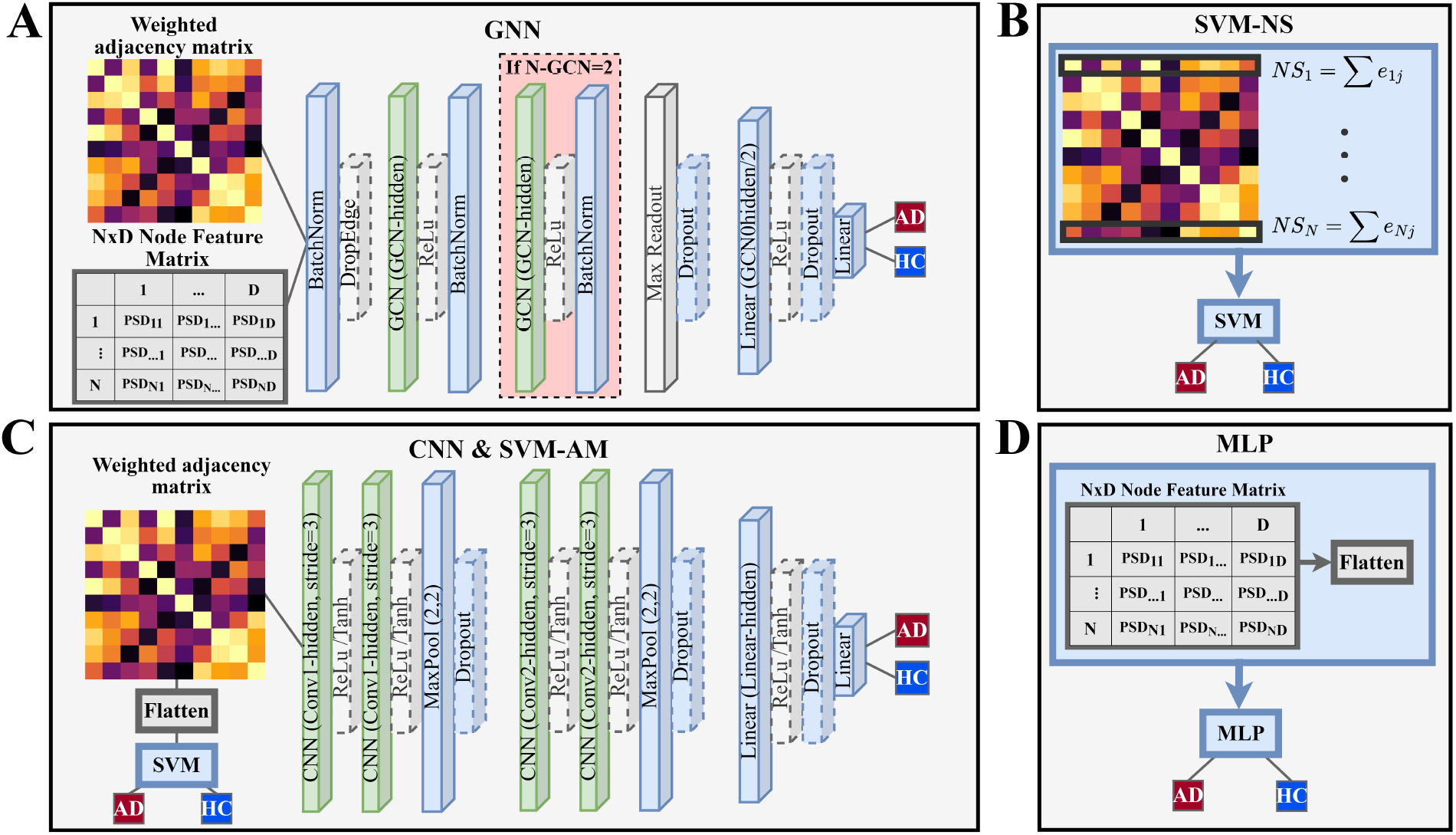
Overview of model architectures developed for classification of AD from EEG-FC-based graphs. (A) A graph neural network (GNN) takes weighted featured brain graphs with *N* nodes represented by a weighted adjacency matrix and a node feature matrix (ℝ^*N×D*^, *D* = 100) where the node features consist of power spectral densities (PSD, 0 – 100*Hz*). The N-GCN hyper-parameter controls the number of graph convolutional layers. (B) Support vector machine trained using the node strengths (i.e. the sum of edge weights of neighbouring nodes) as input features (SVM-NS). (C) Convolutional neural network (CNN) trained on the brain graphs represented by weighted adjacency matrices. Alternatively, the weighted adjacency matrix is flattened and used as input to a support vector machine (SVM-AM). (D) Node feature matrix (ℝ^*N×D*^) with power spectral densities across all EEG channels is used to train a multilayer perceptron (MLP).

## II. DATA AND PRE-PROCESSING

The EEG dataset consists of 20 AD patients and 20 healthy control participants (HC) below 70 years. A subset of this dataset has been previously used in Blackburn et al. [5]. All AD participants were recruited in the Sheffield Teaching Hospital memory clinic. AD participants were diagnosed between one month and two years before data collection, and all were in the mild to moderate stage of the disease at the time of recording. Age and gender-matched HC participants with normal neuropsychological tests and structural MRI scans were recruited. The EEG data used in this study was approved by the Yorkshire and The Humber (Leeds West) Research Ethics Committee (reference number 14/YH/1070). All participants gave their informed written consent.

EEG was acquired using an XLTEK 128-channel headbox, Ag/AgCL electrodes with a sampling frequency of 2 kHz using a modified 10-10 overlapping a 10-20 international electrode placement system with a referential montage with a linked earlobe reference. The recordings lasted 30 minutes, during which the participants were instructed to rest and not to think about anything specific. Within each recording, there were two-minute-long epochs during which the participants had their eyes closed (alternating with equal duration eyes-open epochs, not used in this work).

All the recordings were reviewed by an experienced neurophysiologist on the XLTEK review station with time-locked video recordings (Optima Medical LTD). For each participant, three 12-second-long artefact-free epochs were isolated. Finally, the following 23 bipolar channels were created: F8–F4, F7–F3, F4–C4, F3–C3, F4–FZ, FZ–CZ, F3–FZ, T4–C4, T3–C3, C4–CZ, C3–CZ, CZ–PZ, C4–P4, C3–P3, T4–T6, T3–T5, P4–PZ, P3–PZ, T6–O2, T5–O1, P4–O2, P3–O1 and O1–O2 [5].

### A. EEG Pre-processing

First, a zero-phase 5^th^ order Butterworth filter is employed to remove frequencies below 0.1 Hz and above 100 Hz; a zero-phase 4^th^ order Butterworth stop-band filter is used to remove frequencies between 49 and 51 Hz related to power-noise. The EEG data were then down sampled to 250 Hz using an 8^th^ order Chebyshev type I filter and scaled to zero mean and unit standard deviation.

In order to increase the sample size and to demonstrate that the classification performance is epoch independent, the 12-seconds-long epochs were split into 3-second-long non-overlapping segments. Thus, for each subject, there are 12 EEG segments. Finally, frequency bands are created from each EEG segment using a zero-phase 5^th^ order Butterworth filter. Six frequency bands are considered: *δ* (0.5-*4Hz*), *θ* (4-*7Hz*), α (7-15Hz), *β* (15-31Hz), γ (31 - 100Hz) and full (0.5 - 100Hz).

## III. METHODS

### A. Functional-Connectivity-based Brain Graph Inference

In this paper, we selected eight commonly used methods for constructing brain graphs from EEG signals, namely: the absolute value of Pearson’s correlation (corr), spectral coherence (coh), the imaginary part of coherency (iCOH), phase lagged index (PLI), weighted phase lagged index (wPLI), phase locking value (PLV), mutual information (MI) and amplitude envelope correlation (AEC).

We estimate FC brain graphs for each EEG segment and frequency band separately. Thus, for each subject, we obtain 72 brain graphs (12 segments × 6 frequency bands). A brain graph *G* can be represented by an *N × N* adjacency matrix *A* where *N* = 23. As we consider only FC measures, all edges are undirected, and thus the number of inferred edges can be reduced from *N*^2^ to [*N* × (*N* - 1)/2]. However, for simplicity, we keep the *N*^2^ edges in the *N*× *N* adjacency matrix A. Thus, each entry of the adjacency matrix 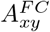 represents the edge weight between nodes, i.e. the dependency of EEG signals 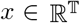 and 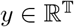 are measured by the connectivity measure *FC* where *T* is the signal length. All of the selected measures are normalised to [0,1] where 0 indicates no coupling and 1 indicates a perfect coupling.

The adjacency matrix using the absolute values of Pearson’s correlation coefficients between nodes *x* and *y* is given by:

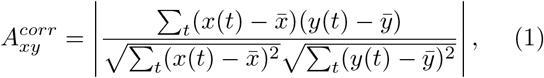

where *x*(*t*) is the value of signal *x* at time *t*, and 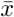 is the mean of *x*. The absolute value is calculated as we are only interested in the coupling magnitude. Next, the adjacency matrix of coherence is given by:

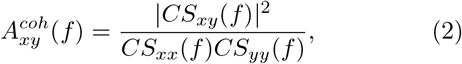

where *CS_xy_* and *CS_xx_* are cross-spectral and auto-spectral densities respectively at frequency *f*. The coherence within a frequency band *B* is then calculated as the mean of 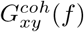 where *f* ∈ *B*.

The imaginary part of coherency (iCOH) measures phase consistency similar to coh and accounts for volume conduction effects. The adjacency matrix using iCOH is computed as:

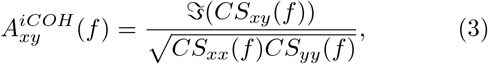

where 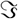 denotes the imaginary component. The iCOH within a frequency band *B* is then calculated as the mean of 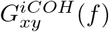 where *f* ∈ *B*

The phase and amplitude of an EEG signal at time *t* can be calculated from the analytic representation *z* of signal *x*

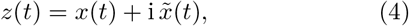

where i is the imaginary component and 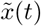 is the corresponding Hilbert transform. Then the phase and amplitude can be obtained from *z*(*t*) as

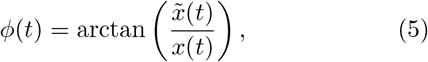

and

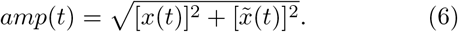

Phase lag index (PLI) quantifies the asymmetry in phase distributions of two signals and measures only nonzero phase locking [27]. The adjacency matrix using PLI is defined as:

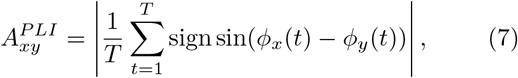

where *ϕ_x_* is obtained using Eq. 5. Weighted phase lag index (wPLI) is an extension of PLI, which aims to remove the effects of amplitude and volume conduction by maximally weighting the ±90 deg phase differences and thus omitting uniformly driven differences [28]. The adjacency matrix using wPLI is computed as

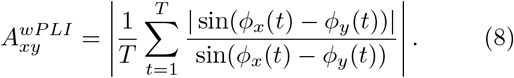

Phase locking value (PLV) is another approach to quantify the consistency of phase differences between signals, and its associated adjacency matrix is computed as

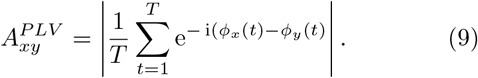

Amplitude envelope correlation (AEC) aims to quantify the coupling based on the amplitudes of the signals. The adjacency matrix using AEC is computed with Eq. 1 where *x* and *y* are the amplitudes of respective signals computed using Eq. 6.

Mutual information (MI) quantifies the amount of known information about a second signal after observing the first signal. The adjacency matrix using MI is calculated as:

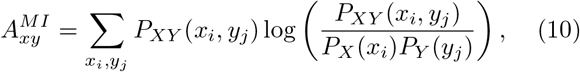

where *P_XY_* and *P_X_* are the joint and marginal probability distributions, respectively.

#### 1. Edge filtering methods

It is worth noting that we did not use any corrections for false positives. Thus, the true brain graph structure might be masked by noise due to spurious coupling. Traditionally, a surrogate threshold might be used to control such spurious edges. However, such a procedure is computationally expensive, as it requires re-computing the connectivity measure on multiple random surrogate versions of the original signals, to estimate a null surrogate distribution. Instead, we implement two edge-filtering methods to select only important edges and thus produce sparse graphs. Compared to the surrogate threshold method, edge-filtering is a fast and efficient, albeit naive method to deal with potentially noisy brain graphs. We also utilise the fully-connected graphs, i.e. without any edge selection, in the classification models in order to test the effect of edge-filtering.

The first edge-filtering method is an FC-strength-based top-k% filter (*k* ∈ {10, 20, 30}), which selects only the top k% strongest edges of the given graph and removes the rest. This approach assumes that edge weight, i.e. the connectivity strength, is directly related to the importance of an edge. However, this assumption might not be valid.

A minimum-spanning-tree-based filter (MST-k), also orthogonal minimum spanning tree [29], addresses this concern as it selects a mix of edge weights and always produces a connected graph, i.e. a path exists among all nodes. Briefly, the MST algorithm [30] aims to extract a backbone of a graph with *N* nodes by selecting *N* – 1 edges, such that the sum of weights is minimised. We use Prim’s algorithm for computing MST [30]. In the case of brain graphs, a stronger edge weight implies a higher degree of coupling; thus, we use an inverted MST algorithm which maximises the sum of weights instead. When *k* = 1, MST-k is equal to a single iteration of the MST algorithm. For *k* > 1, the edges selected by the previous iterations are removed from the graph, and the MST algorithm is re-run. Thus, MST-k filter selects *k*(*N* – 1) edges.

### B. Graph Neural Network Classification

A graph neural network (GNN) is an extension of an artificial neural network that is capable of learning on graph-structured data. Specifically, we implement a graph convolutional network (GCN) for a graph classification task (Fig. 1A).

The input to the GCN classifier is in the form of a graph: *G* = {*N,E,F*}, where *N, E*, and *F* are sets of nodes, edges and node features, respectively. The nodes are fixed in our case, as this is the number of EEG electrodes. The set of edges *E* is given by the adjacency matrix *A* computed by the FC measures introduced in the previous section. Finally, the node feature matrix *F* is an *N* × *D* matrix where each row encodes a *D*-dimensional feature for the corresponding node. Specifically, power spectral density (PSD) is computed over 1 Hz increments in an interval between 0 and 100 Hz, forming a 100-dimensional node feature vector (i.e. *D*=100).

GCN is based on the message-passing framework, which assumes that neighbouring nodes should have similar node features. Briefly, a GCN layer updates the node features (i.e. messages) using the optionally transformed messages collected from neighbouring nodes. On a node level, a single GCN layer effectively aggregates information from the 1-hop neighbourhood of each node. Thus, stacking *L* GCN layers represents aggregation from *L*-hop neighbourhood. Formally, the GCN layer is implemented on a node-level as follows [31]:

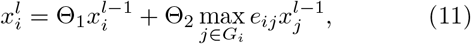

where 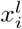 is the node features of node *i* at the *l*^th^ layer, 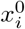 is the *i*^th^ row of the input node feature matrix F, and Θ is a learnable linear transformation, which maps the node features from shape [1, D] to [1, GCN-hidden]. *G_i_* and *e_ij_* are the neighbourhood of node *i* and the edge weight connecting nodes *i* and *j* given by the set of edges *E* respectively. The GCN-hidden is a tunable hyper-parameter of the GCN architecture. A rectified-linear-unit (ReLU) activation is applied to the output of GCN, and batch normalisation is performed [32]. We refer to the nodewise outputs of GCN as node embeddings.

After *L* GCN layers are applied, the output is constructed by node embeddings in the form of a *N* × *H* matrix, where H is the hidden size given by GCN-hidden. In order to produce a graph-level embedding, a maximum readout layer is applied, resulting in an H-dimensional graph embedding *r* for each graph *g*.

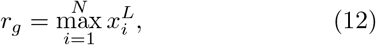

where 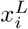 is the output of the *L*^th^ GCN layer for the *i*^th^ node. Following the readout layer, two linear layers are applied to produce the final classification with output dimensions *H*/2 and 2 (number of classes), respectively. Two linear layers were used to allow for further refining of the graph embedding before outputting the predicted class probabilities.

Additionally, in order to improve the generalisability and reduce the risk of overfitting, dropout layers are utilised (1A). Briefly, the dropout layer randomly zeroes elements of the input tensor with p probability drawn from a binomial distribution, where p is a hyperparameter. A dropout is applied to the graph embeddings, i.e. after the readout layer and after the first linear layer. Furthermore, an edge dropout is implemented, which randomly removes edges from the input graph. The inclusion of the edge dropout in the model is controlled by a hyper-parameter.

In summary, the GNN used in this study has several hyper-parameters, as shown in Table I, which control (1) the model architecture, (2) the form of input data, and (3) the training process to prevent overfitting. In particular, (1) is enabled by the number of GCN layers (N-GCN) and the inclusion of edge dropout (DropEdge); (2) is enabled by frequency band and edge filter; and finally, (3) is enabled by dropout probability (drop-p), learning rate, gamma and batch size.

**TABLE I:** Possible values for hyper-parameters of GNN, SVM-NS, SVM-AM, CNN and MLP.

|  | Hyper-parameter | Values | GNN | SVM-NS | SVM-AM | CNN | MLP |
| --- | --- | --- | --- | --- | --- | --- | --- |
| 1 | Cost | [0,1] |  | ✓ | ✓ |  |  |
| 2 | Batch size | {16, 32, 64, 128, 256} | ✓ |  |  | ✓ | ✓ |
| 3 | Learning rate | {0.0001, 0.001, 0.01, 0.1} | ✓ |  |  | ✓ | ✓ |
| 4 | GCN_hidden | {256, 516, 1024, 2048, 4096} | ✓ |  |  |  |  |
| 5 | N_GCN | {1,2} | ✓ |  |  |  |  |
| 6 | N_Linear | {1,2,3,4,5} |  |  |  |  | ✓ |
| 7 | Conv1_hidden | {16, 32, 64, 128} |  |  |  | ✓ |  |
| 8 | Conv2_hidden | {16, 32, 64, 128} |  |  |  | ✓ |  |
| 9 | Linear_hidden | [32, 64, 96, 160, 256, 516] |  |  |  | ✓ | ✓ |
| 10 | Activation | {ReLU, Tanh} |  |  |  | ✓ | ✓ |
| 11 | DropEdge | {True, False} | ✓ |  |  |  |  |
| 12 | Dropout_p | {0.1, 0.25, 0.5, 0.75, 0.8, 0.9} | ✓ |  |  | ✓ | ✓ |
| 13 | Frequency band | delta, theta, alpha, beta, gamma, full | ✓ | ✓ | ✓ | ✓ |  |
| 14 | Edge filter | full, top-{10,20,30}, MST-{1,2,3} | ✓ | ✓ | ✓ | ✓ |  |
| 15 | Gamma | {0.8, 0.85, 0.9, 0.95} | ✓ |  |  | ✓ | ✓ |

### C. Baseline Models

In order to enable a fair assessment of the advantages of using graph-based learning (i.e. the GNN), four baseline classifiers are trained and compared. These baseline models utilise the same graph-structured input data extracted using different FC measures, frequency bands and edge filters, and the same evaluation process. Thus, we argue this to be a fair comparison of models.

The three selected baseline models are based on previously used classifier strategies for learning on FC brain graphs: SVM trained on node strength (SVM-NS) [20], SVM trained on vectorised adjacency matrix (SVM-AM) [21], and CNN trained on image of adjacency matrix (CNN) [22, 33]. Additionally, we train a multilayer perceptron (MLP) on the flattened node feature matrix that was previously used to train the GNN models.

#### 1. Support vector machine baseline models

The SVM-NS and SVM-AM are both trained using an SVM classifier. SVM has only one hyper-parameter, namely the cost, as shown in Table I. Additionally, in order to select an appropriate kernel for SVM, we include two kinds of kernels as hyper-parameters: radial and polynomial (up to 3^rd^ order). Both of the SVM-based baseline models are trained on manually extracted features. All features are first normalised to zero mean and unit standard deviation.

The SVM-NS is trained on node strengths (Fig. 1B). Node strength is defined as the sum of edge weights of one node and can be interpreted as a measure of node importance. Thus, each brain graph is represented by an 23-dimensional feature vector *NS* = (*ns*_1_,*ns*_2_,…,*ns_N_*), where *N* is the number of nodes (*N* = 23).

The SVM-AM is trained on vectorised weighted adjacency matrices (Fig. 1D). As we use only undirected FC measures, the *N* × *N* adjacency matrix of a brain graph is symmetric. Thus, we can use the upper triangular matrix only and flatten it to form a 253-dimensional feature vector (*N* × (*N* - 1)/2). Principal component analysis (PCA) is optionally employed for dimensionality reduction with the number of components selected, such that 95% of the variance is captured. The inclusion of the PCA step is controlled by a hyper-parameter.

#### 2. Convolutional neural network

CNN classifiers are trained on the weighted adjacency matrices of the brain graphs. As the adjacency matrix is a square matrix, it is simple to convert it to an image on which a CNN can be trained.

The CNN architecture consists of two convolutional blocks and a final classifier, as shown in Fig. 1C. Each convolution block contains two convolutional layers (stride = 3), followed by a maximum pooling layer and a dropout layer. The final classifier consists of two linear layers with a dropout between them. We created several hyper-parameters to control the CNN. The number of convolutional filters within each block is controlled by the Conv1 and Conv2 hyper-parameters. Similarly, the hidden size of the linear layers is controlled by the Linear-hidden hyper-parameter. Additionally, there are hyper-parameters controlling the dropout probability, the choice of the activation function (ReLU or Tanh), and the batch size as shown in Table I.

#### 3. Multilayer Perceptron

MLP classifiers are trained using the flattened node feature matrix *F* ∈ ℝ^*N* ×*D*^, where *D* is the PSD computed over the range 1-100 Hz. Thus the entry *F_ij_* corresponds to PSD of the *i*^th^ node at frequency *j*. The MLP is thus trained on the input used to train the GNN models, but without leveraging the topological information provided by the FC graph. The MLP architecture is controlled by the following hyper-parameters: N-Linear (number of layers), Linear-hidden (hidden size). Additionally, there are hyper-parameters controlling the dropout probability, the choice of the activation function (ReLU or Tanh), and the batch size as shown in Table I.

### D. Model evaluation and implementation

The EEG preprocessing, brain graph construction, and model evaluation are implemented in R 4.1.2 [34] using in-house scripts, and caret [35] for SVM training. The training of CNN and GNN classifiers is implemented using PyTorch 1.10 [36] and PyTorch Geometric 2.0.2 [37].

The models are trained and evaluated based on repeated 20-fold cross-validation (CV). A 5 times repeated CV is used in order to identify the best combination of hyper-parameters for all models and FC measures. The folds used for CV are created, such that samples from the same subject are kept within a single fold in order to prevent information leakage. We use a smaller number of repetitions in order to reduce the computational cost of training CNN and GNN models. Hyper-parameter values are selected using random optimisation, where the values of all hyper-parameters are selected randomly. 200 iterations of random optimisation are performed for each combination of FC measure and model type. The hyperparameters of all three model types and their possible values are summarised in Table I.

The best-performing models are selected using the area under the sensitivity-specificity curve (AUC), i.e. one model per each combination of FC measure and model type. In order to assess the stability of the selected models, 50 times repeated CV is performed. The performance errors are computed using the maximum difference between the mean and 5^th^ and 95^th^ quantiles. This approach does not assume a normal distribution and results in conservative error estimates.

The CNN and GNN models are trained using an Adam optimiser with an exponential learning rate decay (controlled by the gamma hyper-parameter) and crossentropy loss function. The models are trained for 300 epochs with an early stopping after 15 epochs if the loss stops decreasing.

## IV. RESULTS

Brain graphs were inferred for each 3-second-long EEG segment by using several commonly used FC measures, which aim to quantify both the linear and nonlinear coupling between pairs of brain signals. The brain graphs were then used as an input to train the GNN brain-graph-classifier. Moreover, four baseline models were trained on these brain graphs in order to demonstrate which type of classifier performs the best. AUC is used to select the best model.

Table II reports the AUC values and the 95% confidence intervals of the SVM-NS, SMV-AM and CNN baseline models and GNN across the 8 FC measures. Note that the MLP baseline is not included here, since it does not utilise the FC brain graphs. Additionally, the performance of the baseline GNN using Euclidean distance between spatial positions of EEG (GNN-euclid) is reported in Table II as well. The hyper-parameter values of the best models from their respective categories are reported in Table IV. The averaged sensitivity-specificity curves of these models are shown in Fig. 2.

**FIG. 2:**
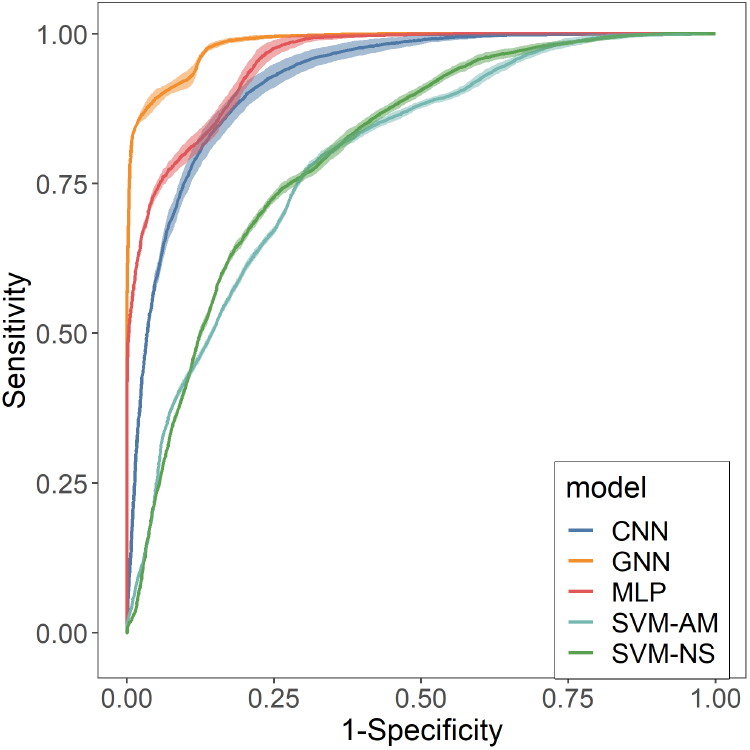
Averaged Sensitivity-Specificity curves of the best models of their respective categories with 95% confidence intervals (ribbon).

**TABLE II:** AUC of GNN, SVM-NS, SVM-AM and CNN models across different FC measures measured by 50-repeated 20-fold cross-validation. The ‘euclid’ entry refers to the baseline GNN model with a fixed graph structure based on the spatial distance of EEG electrodes.

| FC | GNN | SVM-NS | SVM-AM | CNN |
| --- | --- | --- | --- | --- |
| AEC | **0.984 ±0.002** | 0.75 ±0.022 | 0.734 ±0.01 | 0.886 ±0.019 |
| coh | 0.982 ±0.002 | 0.773 ±0.007 | 0.784 ±0.008 | 0.901 ±0.014 |
| corr | 0.982 ±0.003 | 0.764 ±0.008 | 0.773 ±0.008 | 0.887 ±0.014 |
| euclid | 0.978 ±0.004 | **X** | **X** | **X** |
| iCOH | **0.984 ±0.002** | 0.656 ±0.017 | 0.648 ±0.014 | 0.876 ±0.01 |
| MI | 0.981 ±0.002 | **0.807 ±0.008** | 0.779 ±0.013 | **0.924 ±0.015** |
| PLI | 0.982 ±0.005 | 0.761 ±0.057 | 0.603 ±0.036 | 0.893 ±0.015 |
| PLV | 0.982 ±0.003 | 0.766 ±0.008 | **0.793 ±0.007** | 0.9 ±0.014 |
| wPLI | **0.984 ±0.003** | 0.66 ±0.069 | 0.637 ±0.024 | 0.869 ±0.016 |

All baseline models perform worse than all of the GNN models across all FC measures as shown in Table II. Even the best baseline model, MLP (AUC=0.95), achieves lower performance than the worst GNN model, GNN-euclid (AUC=0.978).

From Table II, we can also see that the GNN models trained using FC-based brain graphs perform better than GNN-euclid, which was trained using a static graph structure.

Furthermore, we report the effect of frequency bands and edge filtering methods on the performance of the trained models in the supplementary materials. Figure S3 and tables S1-S3 report these effects of frequency bands. Figure S4 and Tables S4-S6 report these effects of edge filtering methods.

## V. DISCUSSION

We trained GNN models over several commonly used FC measures. For comparison, we trained four baseline models. The results suggest that the GNN outperforms all baseline models across all FC measures (Table II). Moreover, neural-network-based models (GNN, CNN and MLP), which perform automatic feature extraction, perform decisively better than the classical machine learning approaches (SVM-AM and SVM-NS) that rely on manually engineered features.

We argue that the relatively low performance of the machine learning approaches is caused by the inability to remove noise-contaminated information from the input features. This is likely exacerbated by the lack of false positives control during the brain graph inference, which would limit the number of edges caused by spurious coupling. We suggest that the neural network-based models can solve this issue by using weight regularisation and dropout layers, designed to learn generalisable features insensitive to noise.

It could be argued that the GNN models perform better than CNN and MLP, because they are trained using two input information sources, i.e. the FC weighted brain graph and the node feature matrix with power spectral density. This is a unique property of GNN as it can aggregate information from both inputs. Moreover, to the best of our knowledge, GNN is the only model architecture that can process these two inputs simultaneously.

The CNN and MLP baseline models offer an interesting comparison to the GNN, since each is trained using one of the two input information sources. The CNN and MLP baselines show the individual predictive power of the FC-based brain graph and node feature matrix, respectively. The results suggest that the node feature matrix provides a slightly better source of information in the classification task (Table III). However, GNN performs significantly better, and we argue that the comparison with the CNN and MLP baselines highlights the power of GNN in brain-graph classification.

The relatively poor performance of CNN also demonstrates the shortcomings of treating the adjacency matrix of a brain graph as an image. Each pixel of an image has an equal number of neighbouring pixels, and the content of the image depends on the specific spatial ordering of its pixels. Therefore, convolution can be applied to patches of pixels to extract features automatically. This assumption is invalid for a graph where each node can be connected to an arbitrary number of neighbours, and no meaningful ordering of nodes exists. In contrast, graph convolution generalises the convolution to efficiently solve this issue by utilising order invariant operations to aggregate information from neighbouring nodes.

Moreover, the hyper-parameter optimisation has identified a GNN model with two graph convolutional layers as the optimal GNN architecture (Table IV). This means that the GNN aggregates information not only from the nodes connected by an edge directly (i.e., the 1-hop neighbours), but also from the 2-hop neighbours. This suggests the importance of global graph properties in diagnosing AD accurately, in addition to the local properties, which could likely be learned with a single layer. This is in line with the reported loss of small-world properties of AD brain graphs [11–14].

Next, the results demonstrate that the FC-based GNNs also outperform the GNN-euclid model, which utilises a static graph structure (Table II). This suggests that it is preferable to utilise FC-based brain graphs rather than the distance-based static graphs previously used for EEG-GNN tasks [23, 24]. However, it seems that no FC measure offers clearly superior performance compared to the others. Thus, we suggest that future studies need to carefully consider which FC measure to use based on the type of brain coupling they might wish to focus on. However, we do not claim that the brain graphs inferred from various FC measures are necessarily similar from a graph-theoretic perspective. This is supported by the performance differences of the baseline models where some of the FC measures, such as MI, perform consistently well.

Surprisingly, the GNN-euclid model achieves relatively high accuracy despite utilising a fixed graph structure (Table III). The Euclidean brain-graph structure highlights the spatially local relationships between the EEG channels. In contrast, long-range edges have only a low weight. Therefore, we argue that the Euclidean brain graph biases the GNN model to learn local graph features predominantly. On the other hand, the FC-based brain graphs may contain both local and long-range relationships. Previous research suggests that AD-related differences are observed in long-range pathways and global graph properties [10, 11, 13]. In our opinion, the FC-based GNNs outperform GNN-euclid, since they can better capture both the local and global differences on the graph level.

**TABLE III:** Detailed performance metrics of best performing models (selected based on AUC) of each model type.

|  | AUC | Accuracy | Sensitivity | Specificity |
| --- | --- | --- | --- | --- |
| GNN (AEC) | 0.984 ±0.002 | 91.996% ±0.41 | 97.366% ±0.941 | 86.716% ±1.013 |
| GNN-euclid | 0.978 ±0.004 | 91.147% ±1.278 | 93.678% ±2.46 | 88.658% ±3.497 |
| CNN (MI) | 0.924 ±0.015 | 84.689% ±2.233 | 86.212% ±3.6 | 83.192% ±4.613 |
| SVM-NS (MI) | 0.807 ±0.008 | 73.924% ±0.852 | 73.494% ±0.926 | 74.367% ±0.934 |
| SVM-AM (PLV) | 0.793 ±0.007 | 72.929% ±0.6 | 72.123% ±0.673 | 73.799% ±1.064 |
| MLP | 0.953 ±0.007 | 84.97% ±1.93 | 82.77% ±2.31 | 87.13% ±2.88 |

To further investigate the differences between FC measures on the graph level, we compute an average adjacency matrix for each FC measure across both groups and frequency bands (Figure S1). In Figure 3, we show these matrices for *α* and *θ* frequency bands as these are utilised by the best performing models (Table IV). The brain graphs are relatively similar across the FC measures. In the θ band, increased connectivity can be observed in AD compared to HC. In contrast, the connectivity seems to be decreased in AD in the α band. These differences are well documented in the literature [8–10].

**FIG. 3:**
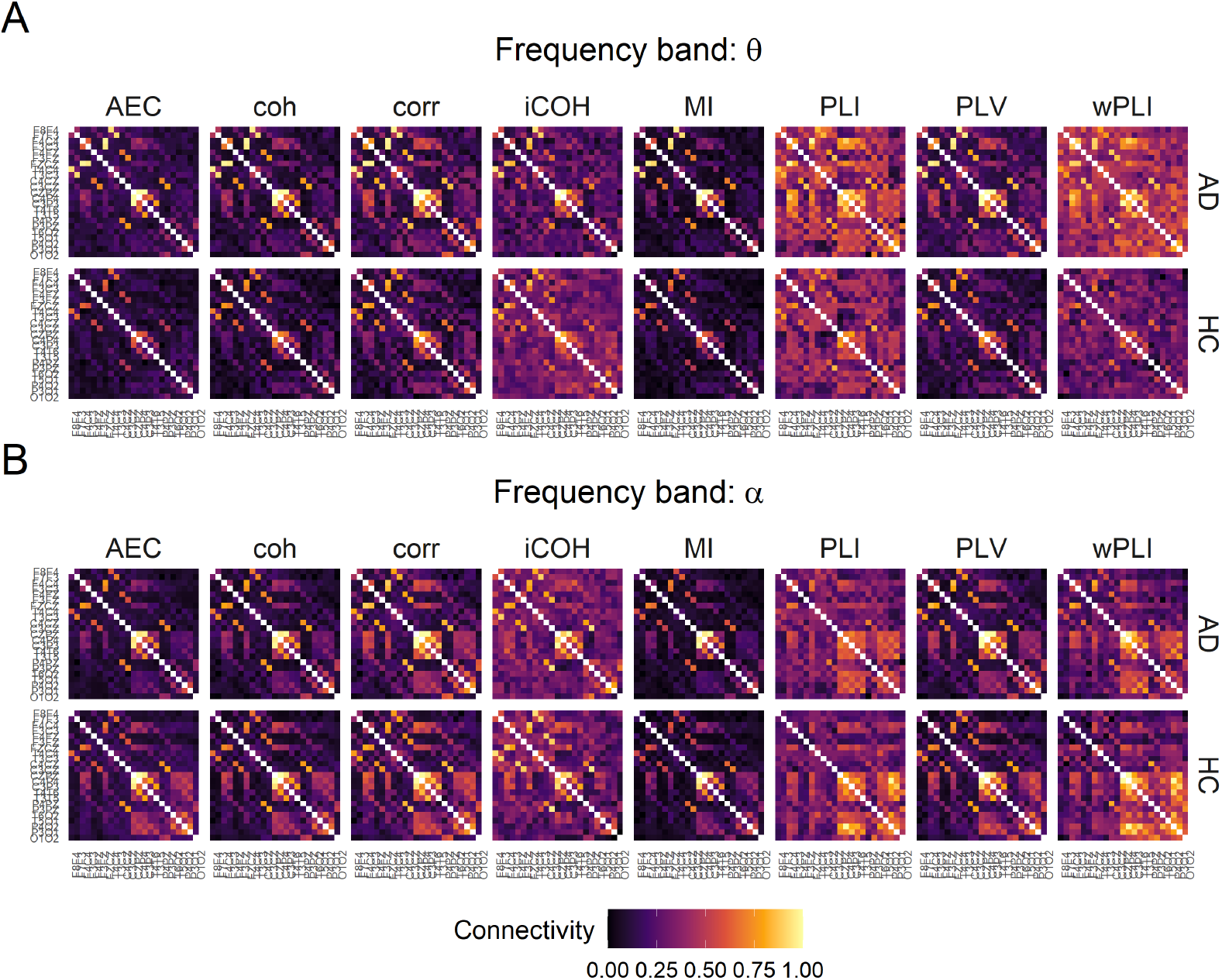
Averaged adjacency matrices of AD and HC cases measured with various functional connectivity measures in A) *θ* (best in CNN, SVM-NS and SVM-AM models), and B) *α* (best in GNN model) frequency bands.

**TABLE IV:** Hyper-parameter values of the best performing GNN, SVM-NS, SVM-AM, CNN and MLP measured by AUC.

| Hyper-parameter | GNN | SVM-NS | SVM-AM | CNN | MLP |
| --- | --- | --- | --- | --- | --- |
| FC measure | AEC | MI | PLV | MI |  |
| Cost |  | 0.5 | 0.5 |  |  |
| Batch size | 32 |  |  | 64 | 32 |
| Learning rate | 0.001 |  |  | 0.001 | 0.01 |
| GCN_hidden | 1024 |  |  |  |  |
| N_GCN | 2 |  |  |  |  |
| N_Linear |  |  |  |  | 4 |
| Conv1_hidden |  |  |  | 128 |  |
| Conv2_hidden |  |  |  | 64 |  |
| Linear_hidden |  |  |  | 96 | 4096 |
| Activation/Kernel |  | Radial | Poly (1<sup>st</sup>) | ReLU | ReLU |
| DropEdge | TRUE |  |  |  |  |
| Drop_rate | 0.9 |  |  | 0.1 | 0.05 |
| Frequency band | alpha | theta | theta | theta |  |
| Edge filter | Top-20% | MST-3 | full | full |  |
| Gamma | 0.9 |  |  | 0.9 | 0.95 |

Interestingly, all FC measures detect a well-defined cluster containing mostly parietal and occipital EEG channels. The strength of this cluster distinguishes the AD from HC consistently across FC measures. We speculate that this cluster contributes most of the predictive information for the classification models. However, since the GNN architecture is a block-box model, it would be difficult to confirm our speculation.

Next, the optimised model architectures suggest that using edge-filtering and filtering the EEG signal within a frequency band improve the model performance. A detailed report of differences between the edge-filtering methods and frequency bands across the FC measures and model types is included in the supplements (Figures S3-S4 and Tables S1-S6). Briefly, the best GNN model utilises the α frequency band, and CNN, SVM-AM and SVM-NS utilise the θ frequency band (Table IV), suggesting that frequency-centred brain graphs should be preferred over the full-frequency-range brain graphs. The selection of these frequency bands is not surprising, as they are both well known to be altered in patients with AD [8–10]. In contrast, the effect of edge-filtering is not so apparent as only the GNN and SVM-NS models use edge-filtering with top-20% and MST-3, respectively. On the other hand, CNN and SVM-AM use unfiltered brain graphs. We expect that a sparse graph is preferable for GNN since there are fewer messages to aggregate while updating the node embeddings. These messages are also less likely to be a product of false-positive brain interaction, thus leading to better node and graph embeddings.

Furthermore, it is worth noting that although GNN accepts two inputs, the relative contribution of each input information is largely unclear. The results suggest that the node feature matrix should contribute slightly more, since the MLP baseline outperforms the CNN (Table III). It could be argued that the GNN uses only the topological information provided by the graph structure to enable message-passing, but the FC is not fully reflected in the node embeddings and graph embeddings by extension. Nevertheless, we believe that the FC information is utilised to some extent by the GNNs since these models perform better than the GNN-euclid, which arguably utilises merely the topological information (Table II). However, the extent to which the information provided by the FC measures is contained within the learned graph embedding remains unclear. One can merely speculate without introducing an additional mechanism into the GNN architecture, which is beyond the scope of this paper.

Finally, the GNN architecture utilised in this study is relatively simple as one of the simplest GCNs was used, and the readout layer merely computes the maximum of the node embeddings. Previous EEG-GNN applications demonstrated the advantages of using more complex graph convolutional layers and edge pooling mechanisms [23]. We hypothesise that exploiting a learnable edge-filtering mechanism akin to that utilised by Zhong et al. [26] might improve the classification compared to the edge-filtering methods used in this study.

## VI. CONCLUSION

GNN is an effective model for learning on graph-structured data, such as FC-EEG brain graphs. However, in the absence of consent about the ideal FC measure for estimating EEG brain graphs, the effect of an FC measure on the performance of GNN classifiers is unclear. In this paper, we have selected eight common FC measures to investigate this effect.

First, we demonstrated that GNN models are superior to classical machine learning and CNN models for brain graph classification. Unfortunately, the utilised GNN architecture is a black-box model. Thus, future work should focus on implementing interpretable GNN architectures that achieve similar performance but additionally offer interpretability, such as which nodes, i.e. brain regions, drive the prediction. Besides providing an opportunity for experts to validate such models, interpretable predictions might also serve in the development of GNN-informed targeted treatment.

Finally, we showed that utilising FC measures to define the brain graph results in improved performance of GNN models compared to a fixed graph structure (i.e. the Euclidean distance between EEG electrodes). While using an FC measure improves the performance, no concrete FC measure can be recommended as the ideal choice. Thus, in future research, the choice of suitable FC measure should be carefully evaluated in the context of the given research question. Alternatively, focusing on fusion methods might lead to developing a novel composite measure of FC.

## ACKNOWLEDGEMENT

The EEG data used in this study was approved by the Yorkshire and The Humber (Leeds West) Research Ethics Committee (reference number 14/YH/1070). All participants gave their informed written consent. Dominik Klepl was supported by A*STAR Research Attachment Programme (ARAP). The work was supported by A*STAR, AI, Analytics and Informatics (AI3) Horizontal Technology Programme Office (HTPO) seed grant C211118015.

